# Aboave-Weaire’s law in epithelia results from an angle constraint in contiguous polygonal lattices

**DOI:** 10.1101/591461

**Authors:** Roman Vetter, Marco Kokic, Harold Gómez, Leonie Hodel, Bruno Gjeta, Antonella Iannini, Gema Villa-Fombuena, Fernando Casares, Dagmar Iber

## Abstract

It has long been noted that the cell arrangements in epithelia, regardless of their origin, exhibit some striking regularities: first, the average number of cell neighbours at the apical side is (close to) six. Second, the average apical cell area is linearly related to the number of neighbours, such that cells with larger apical area have on average more neighbours, a relation termed Lewis’ law. Third, Aboav-Weaire’s (AW) law relates the number of neighbours that a cell has to that of its direct neighbours. While the first rule can be explained with topological constraints in contiguous polygonal lattices, and the second rule (Lewis’ law) with the minimisation of the lateral contact surface energy, the driving forces behind the AW law have remained elusive. We now show that also the AW law emerges to minimise the lateral contact surface energy in polygonal lattices by driving cells to the most regular polygonal shape, but while Lewis’ law regulates the side lengths, the AW law controls the angles. We conclude that global apical epithelial organization is the result of energy minimisation under topological constraints.

## INTRODUCTION

Epithelia are polarized tissues. This means that cell properties change along the apical-basal axis (Fig. 1a). In particular, cells adhere tightly on the apical side via an adhesion belt composed of Cadherins, while they bind to the extracellular matrix on the basal side. When viewed from the apical side, the tissue appears as a contiguous polygonal lattice (Fig. 1b). The organisation of the polygonal lattice may appear random at first sight, but it has previously been noted to follow certain phenomenological laws. First, even though the fraction of cells with a certain number of neighbours differs hugely between epithelial tissues (Fig. 1c), the average number of neighbours, 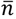, is always close to six (Fig. 1d) [1–7] because of topological constraints in 2D contiguous polygonal lattices [4, 8]. In fact, in the limit of an infinite number of polygons, the average number of neighbours is exactly

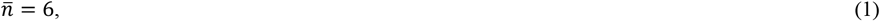

if three cells meet at each vertex; this can be derived from Euler’s formula, *v*−*e*+*f*=2, which formulates a relationship between the number of vertices, *v*, edges, *e*, and faces, *f*. Second, the average apical area is linearly related to the number of cell neighbours, *n*, a relation termed Lewis’ law (Fig. 1e), i.e.

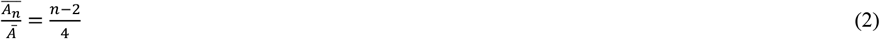

**Figure 1:**
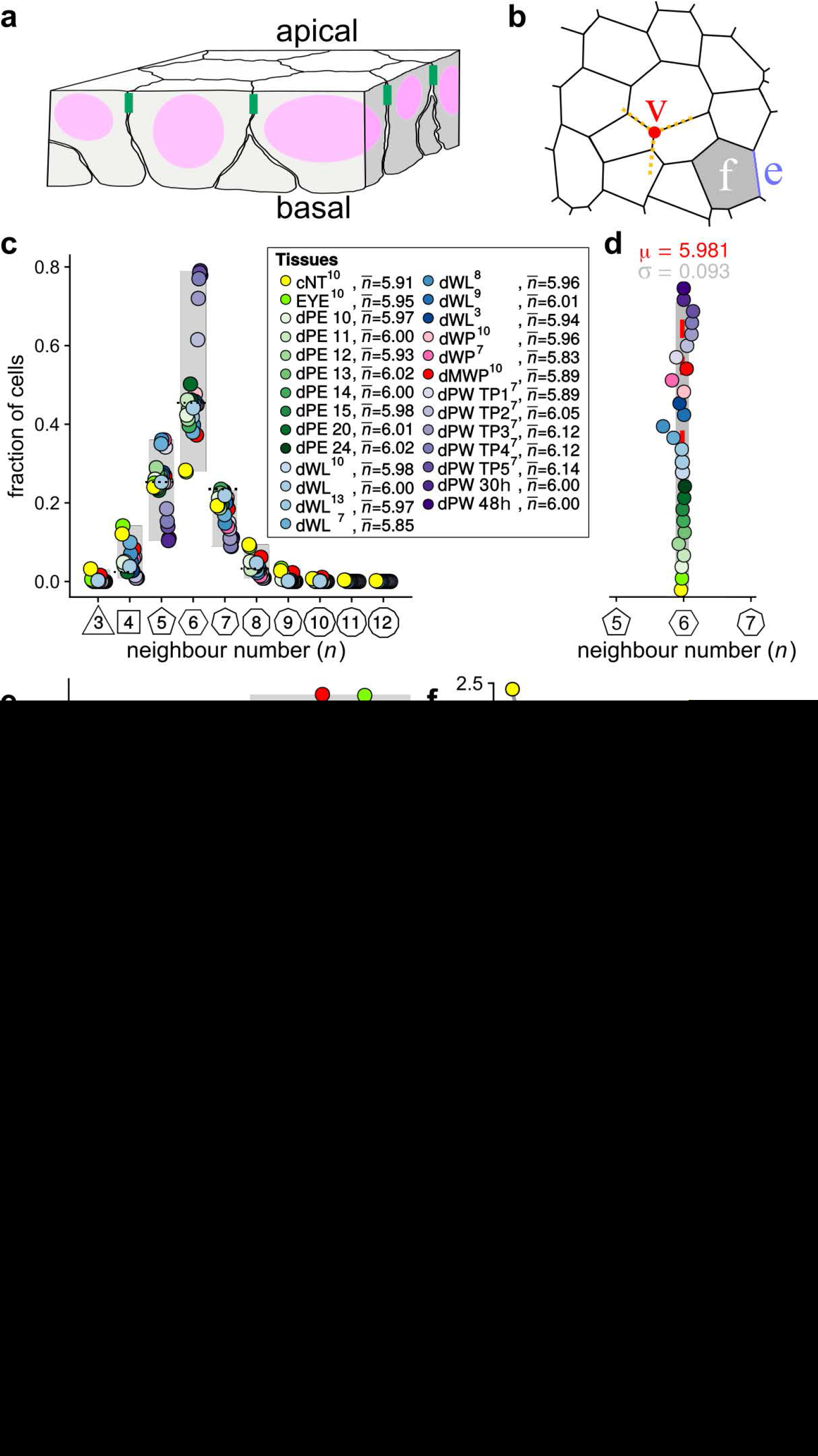
Principles of Epithelial Organisation. **(a)** A cartoon of an epithelial tissue. The tissue is polarized with an apical and basal side. At the apical side cells adhere tightly via adhesion junctions (green). Nuclei are depicted in pink. **(b)** The epithelial cells (denoted as faces - *f*) form a contiguous polygonal lattice on the apical side. Typically, three edges, *e*, meet in a vertex, *v*. **(c)** Tissues differ widely in the frequency of neighbour numbers. The legend provides the measured average number of cell neighbours for each tissue and the references to the primary data [1–7]. Data points for *n* < 3 were removed as they must present segmentation artefacts. The tissue samples are: Chick neural tube epithelium (cNT) [2], *Drosophila* eye disc (EYE) [2], *Drosophila* peripodal membrane from the larval eye disc (dPE), *Drosophila* larval wing disc (dWL) [1–6, 19] (Kokic et al, submitted), *Drosophila* pre-pupal wing disc (dWP) [1, 2], *Drosophila* mutant wing pre-pupa (dMWP: the expression of myosin II was reduced in the wing disc epithelium by expressing an UAS-zipper-RNAi using the C765-Gal4 line) [2], and *Drosophila* pupal wing disc (dPW) [1]. In case of the *Drosophila* peripodal membrane from the larval eye disc (dPE), the numbers indicate the developmental age as the number of ommatidial rows that have formed. In case of the *Drosophila* pupal wing disc (dPW) [2], TPx indicates subsequent, but not further specified pupal time points [1]; 30h and 48h specifies the time post puparium formation. **(d)** The measured average number of cell neighbours is close to the topological requirement 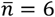 in all tissues; see panel c for the colour code. **(e)** The relative average cell area, 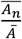, is linearly related to the number of neighbours, *n*, and follows Lewis’ law (Eq. 2, black line) in all reported tissues. The colour code is as in panel c, but data is available only for a subset of those tissues. Few cells with more than 7 neighbours have been measured and estimates of the mean areas are therefore unreliable (shaded part). **(f)** The normalised side length of a regular polygon with *n* vertices for a fixed area. **(g)** The AW law formulates a relationship between the number of neighbours, *n*, that a cell has and the average number of neighbours, *m_n_*, that its direct neighbours have. The product *m_n_* · *n* can be determined by summing over all *n* neighbours. **(h)** The epithelial tissues follow the AW law (black line). The colour code is as in panel c, but data is available only for a subset of those tissues. Panels a-f are reproduced with modifications from (Kokic et al, submitted).

Here, 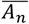 refers to the average apical area of cells with *n* neighbours, and *Ā* refers to the average apical area of all cells in the tissue. We have recently shown that Lewis’ law can be explained with a minimisation of the lateral cell-cell contact surface energy (Kokic et al, submitted). The lateral contact surface energy is minimal if the cell-cell contact area, and thus the combined cell perimeter is minimised. For a given apical area, the cell perimeter is minimal if the polygons are the most regular. To form a contiguous polygonal lattice from regular polygons, these need to all have the same side/edge length. However, for the same polygon area, side lengths differ between polygon types (Fig. 1f). In epithelial tissues, apical areas vary as a result of several processes, most prominently growth and cell division. By following the relationship between polygon area and polygon type as stipulated by Lewis’ law (Eq. 2), the difference in side lengths is minimised between cells. However, equal side lengths are only achieved if the areas followed a quadratic rather than a linear relationship,

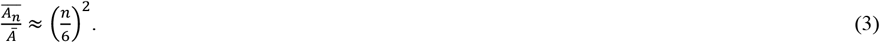

This quadratic relationship only emerges in case of a larger variability in apical areas than previously observed in epithelia. We confirmed this prediction by increasing the area variability in the *Drosophila* larval wing disc, which led to the predicted quadratic relationship (Kokic et al, submitted).

Finally, Aboav-Weaire’s law (AW law) formulates a relationship between the number of neighbours, *n*, that a cell has and the average number of neighbours, *m_n_*, of its direct neighbours (Fig. 1g). Aboav [9] empirically observed for grains in a polycrystal that

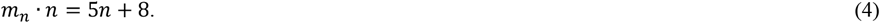

Eq. 4 has previously been found to be a close approximation to the neighbour relationships in epithelial tissue in both plants [10–12] and animals [7] (Fig. 1h). However, Eq. 4 cannot explain regular hexagonal packings, where 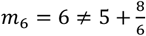. Weaire suggested a refinement of the form

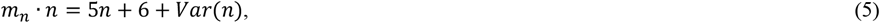

where 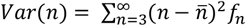 is the variance of the polygon distribution, and *f_n_* is the fraction of cells with *n* neighbours [13]. Eq. 5 applies to hexagonal lattices as *Var*(*n*) = 0, and reduces to Eq. 4 for *Var*(*n*) = 2. Efforts to explain the mathematical basis of Eqs. 4 and 5 have led to a wide range of alternative formulations [14], but no physical or mathematical argument has so far been established to explain the experimental observations.

In the following, we will show that the emergence of the AW law in epithelia can be explained with the minimisation of the lateral contact surface energy. We previously showed that Lewis’ law ensures that the side lengths of the different polygon types are the most similar, so that the different polygon types can fit within a polygonal lattice as the most regular polygons. To form a regular polygon also the internal angles of the polygon must match the polygon type, and to form a contiguous lattice, the internal angles around the shared vertex of adjacent cells must add to 360°. We show that this second constraint results in the AW law.

## RESULTS

### The emergence of the AW law can be explained with the minimisation of lateral surface energy

In the first step, we sought to test whether the AW law could be explained with the minimisation of the lateral contact surface energy. We have previously introduced the simulation framework `LBIBCell` [15] to simulate epithelial tissues (Kokic et al, submitted). `LBIBCell` represents tissues in a 2D plane, and can thus be used to simulate the apical tissue dynamics (Fig. 2a). The simulation framework offers high spatial resolution, i.e., it represents the boundaries of all cells separately such that cells can unbind and the spatial resolution is sufficiently high that cell edges can be curved. Cortical tension as well as cell-cell adhesion are implemented via springs between vertices. The fluid inside and outside the cells is represented explicitly and the fluid dynamics are approximated by the Lattice Boltzmann method. The interaction between the elastic cell boundaries and the fluid is realised via an immersed boundary condition.

**Figure 2:**
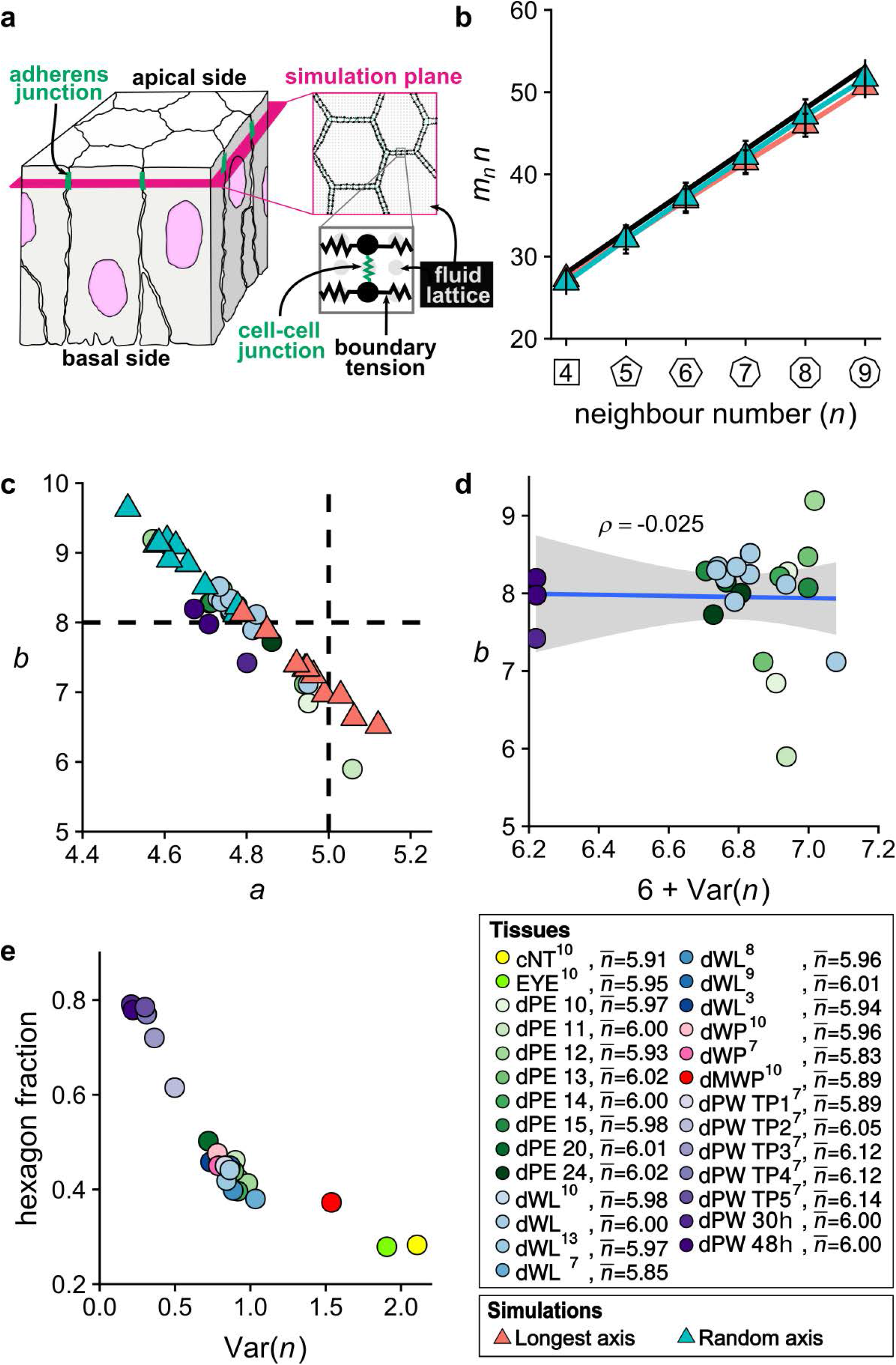
The AW law emerges in LBIBCell Simulations. **(a)** Representation of the apical epithelial surface as a 2D simulation plane in LBIBCell. The cartoon shows a cubical section of a simple columnar pseudostratified epithelium. LBIBCell simulates the tissue mechanics and morphodynamics of the 2D plane (indicated in pink) that goes through the adherens junctions (marked in green), which are located below the apical surface. Cell boundaries are geometrically highly resolved and cell boundaries and cell-cell junctions are modelled with elastic springs. The cartoon and legend are reproduced with modifications from (Kokic et al, submitted). **(b)** The AW law plotted for `LBIBCell` simulations with random cell division (blue) or cell division perpendicular to the longest axis (red). **(c)** The estimated coefficients *a, b* when fitting *m_n_* · *n* = *a* · *n* + *b* to the data (Fig. 1h) and the `LBIBCell` simulations (Fig. 2b). **(d)** The estimated coefficient *b* when fitting *m_n_* · *n* = *a* · *n* + *b* does not positively correlate with 6 + *Var*(*n*) in epithelia. *ρ* is the Pearson correlation coefficient. **(e)** The hexagon fraction versus the variance in the polygon distribution *Var*(*n*) for all available data.

The biological parameters in the simulation are the growth rate, the cell division threshold (i.e. the apical cell area, at which the cell divides), as well as the spring constants representing cortical tension and cell-cell adhesion. We have previously adjusted these to recapitulate the quantitative data from the *Drosophila* larval wing disc, a model epithelial tissue, and we have re-used the same setup as described before (Kokic et al, submitted; Supplementary Material, Table S1). When we simulate the *Drosophila* larval wing disc dynamics while dividing cells either perpendicular to their longest axis (Hertwig’s rule [16]) (Fig. 2b, orange line) or randomly (Fig. 2b, cyan line), the resulting lattice results in polygonal relationships that are close to the AW law (Fig. 2b, black line). The deviation of the simulations from the AW law is similar to that observed for epithelial tissues (Fig. 2c). Thus, if we determine the parameters *a* and *b* for which

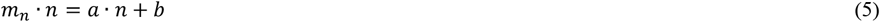

best fits the simulations and the data, the inferred values differ in a similar way from those of the AW law (*a*=5, *b*=8) [9]. We note that the *b* parameter deviates more than the *a* parameter. Weaire linked the *b* parameter to the variance in the observed polygon distribution, i.e. *b* = 6 + *Var*(*n*) (Eq. 5) [13]. However, 6 + *Var*(*n*) does not correlate with the parameter *b* that we infer from the data (Fig. 2d), thus ruling out such a relationship. Given the definition of variance, the hexagon fraction declines with the variance of the polygon distribution (Fig. 2e), and *Var*(*n*) = 2 and thus *b* = 8 as in Aboav’s formulation is observed only for tissues with low hexagon fraction (~30%), which are rare (Fig. 1c).

We conclude that the `LBIBCell` simulations generate a distribution of cell-cell contacts that lead to similar parameters for the AW law as observed in epithelia, but that differ from those specified by Aboav and Weaire. This suggests that the AW law could be explained with a minimisation of the lateral contact surface energy. But why would energy minimisation result in neighbour arrangements that follow the AW law?

### Perfect polygonal lattices do not follow Aboav’s law, but match Weaire’s law

To understand the underlying constraint that leads to the particular form of the AW law, we consider perfect polygonal lattices made of regular polygons. The simplest case is a perfect hexagonal lattice (Fig. 3a, red). Here, all polygons must have the same area. To generate other regular lattices, cells with the appropriate areas must be placed in the correct relative position within the polygonal lattice. The relative areas of the different polygon types must follow the quadratic law (Eq. 3) so that all polygons in the lattice have the same side length. In addition, the internal angles of the cells that meet at a vertex must add up to 360° (Fig. 3b). The internal angles of a regular polygon with *n* neighbours (Fig. 3c) are given by

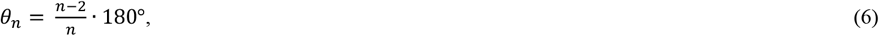

and only in few regular configurations, the angles at each vertex will add up to 360° (Fig. 3a). Consider for instance the case of a lattice made of repetitive elements with one square and two octagons: the 90° angle of a square and the 135° angles of two regular octagons (Fig. 3c) add up to 360° at each vertex (Fig. 3a). If in addition, the octagons have about four times the area of the square (Eq. 3), a regular lattice emerges (Fig. 3a, blue). On the contrary, if the areas of neighbouring cells do not match such requirements, irregular lattices emerge (Fig. 3d).

**Figure 3:**
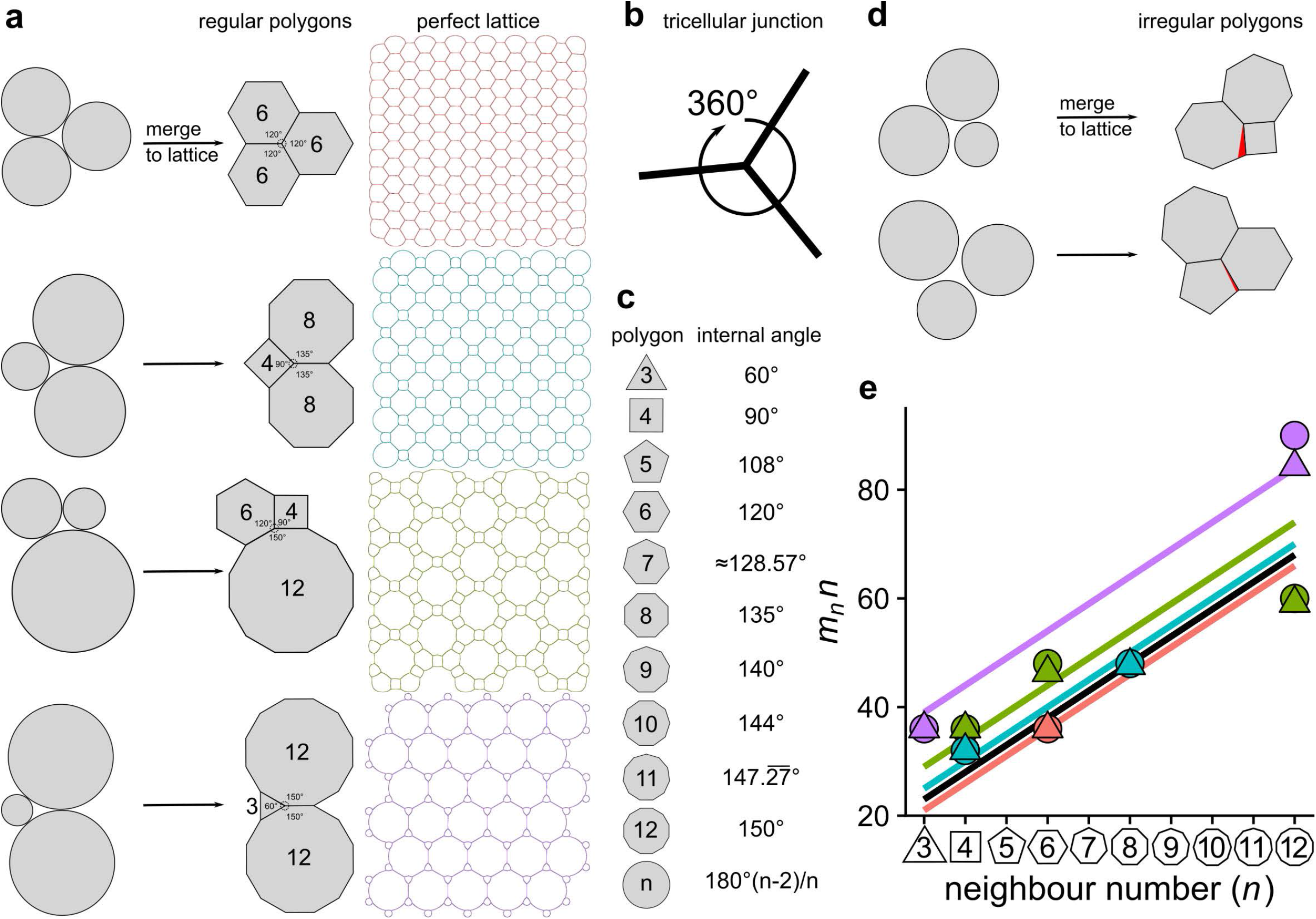
The AW law does not emerge in perfect lattices. **(a)** In case of appropriate relative sizes and positions, adherence of circles results in perfect lattices (grey), as confirmed in `LBIBCell` simulations. **(b)** The angles at each tricellular junction have to add up to 360°. **(c)** Interior angles of regular polygons. **(d)** In case of inappropriate relative sizes and positions, adherence between circles results in imperfect lattices. **(e)** The products of *m_n_* · *n* versus neighbour number *n* for the perfect lattices in panel a deviate from Aboav’s law (Eq. 4, black line), but match Weaire’s law (Eq. 5, coloured lines; colour code as in panel a) rather well. Results from `LBIBCell` simulations (triangles) are close to analytic results (circles).

As noted above, in a hexagonal lattice the average number of neighbours is six, i.e., 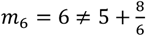 (Fig. 3e, red triangle), which violates Aboav’s law (Eq. 4, Fig. 3e, black line), but matches Weaire’s law (Eq. 5, Fig. 3e, red line). Also, the other arrangements that result in perfect polygonal lattices (Fig. 3a) deviate from Aboav’s law (Eq. 4, black line), but match Weaire’s law (Eq. 5, coloured lines) rather well (Fig. 3e). These perfect lattices also all emerge in `LBIBCell` tissue simulations when adhesive circles with the appropriate relative areas are placed at the appropriate relative positions in a grid (Fig. 3a, coloured lattices) and thus represent the configuration with the lowest lateral surface energy. This shows that not all configurations that minimise the lateral contact surface area follow Aboav’s law.

In summary, regular polygonal lattices follow Weaire’s law more closely than Aboav’s law (Fig. 3e), while epithelial tissues follow Aboav’s law more closely than Weaire’s law (Fig. 2c,d). What is special then about epithelial tissues, and what is the underlying physical principle that leads to these regularities?

### The AW law in epithelial tissues

In an epithelial tissue, the apical areas are predetermined by several active processes, including cell growth, cell division, and interkinetic nuclear migration. As a result, the position of cells with a certain area does not follow a pattern that would allow the emergence of a contiguous lattice made from regular polygons. But, why would these lattices follow the AW law?

According to Euler’s formula, cells have on average six neighbours [4, 8]. The plot of *m_n_* · *n* with *m_n_* = 6 versus *n* is reasonably close to the AW law (Fig. 4a, black line), but clearly deviates (Fig. 4a, green line). In a contiguous lattice, cells have to meet a further constraint, in that at each vertex point, the combined angle must be 360° (Fig. 3b). At each of the *n* vertices, the neighbouring cells should have a combined angle of 360° - *θ_n_* (Fig. 4b, inset). The average angle, 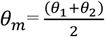, of the two neighbouring cells therefore follows from 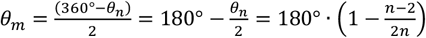, where we used Eq. 6 for *θ_n_*. We then obtain for the average number of neighbours

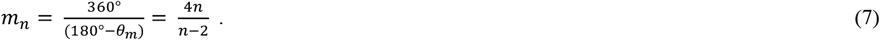

**Figure 4:**
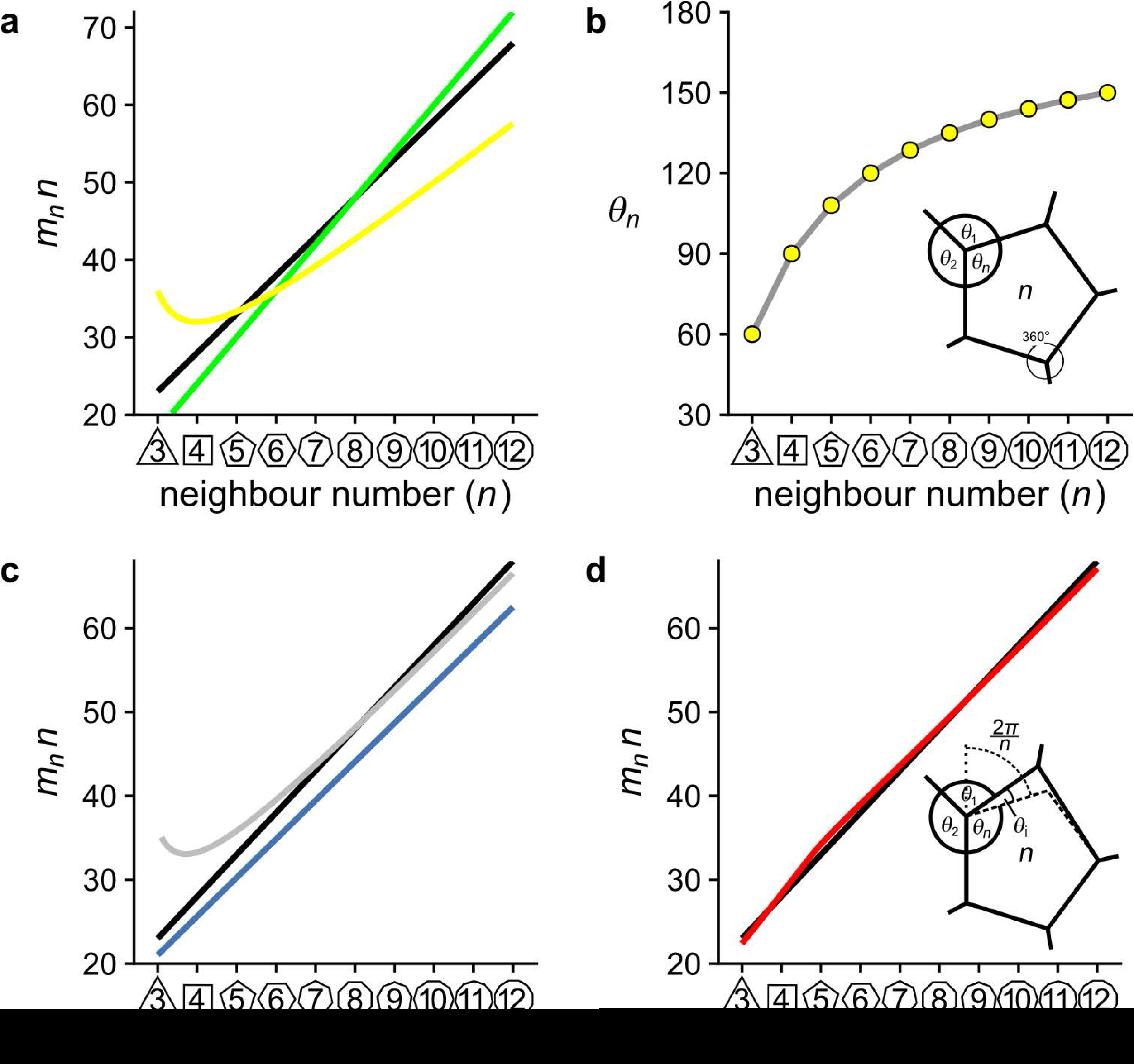
The AW law in epithelial tissues arises because of an angle constraint. **(a)** The product of *m_n_* · *n* versus neighbour number *n* for the AW law (black), *m_n_* = 6 (green), 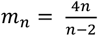 (yellow). **(b)** Interpolation between the angles of regular polygons, 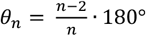, versus the number of vertices *n. Inset*: A cartoon of the angles at a tricellular junction between a cell with *n* neighbours and its two neighbours with angles *θ*_1_ and *θ*_2_. **(c)** Comparison of an analytical approximation for regular polygons (Eq. 8, grey) and its asymptote (*a* ≈ 4.6 and *b* = 7.2) (blue) to Aboav–Weaire’s law (black). **(d)** Comparison of an analytical approximation for irregular polygons (red) to Aboav–Weaire’s law (black). *Inset*: A cartoon of the angles at a tricellular junction in case of an irregular central polygon with angle *θ_n_* + *θ*_i_.

This relationship approximates the AW law for neighbour numbers close to *n* = 5 rather well, but deviates for larger and smaller neighbour numbers (Fig. 4a, yellow line). We note that the internal angle of regular polygons, *θ_n_*, depends non-linearly on the polygon type *n* (Eq. 6, Fig. 4b). As a consequence, two cells, whose angles at a vertex add up to 2*θ_m_* yield a different *m_n_* depending on whether their angles are very similar or very different. If we take this non-linear effect into account (see Supplementary Information for details), and integrate within sensible physiological limits, i.e. *θ*_min_ = 60°, the smallest internal angle of a regular polygon (triangle), and 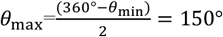, the internal angle of a regular dodecagon (12-gon), we obtain

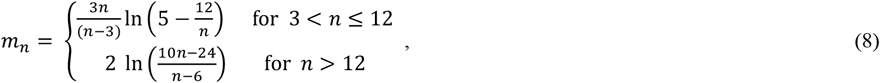

which yields a rather close fit to the AW law for larger *n* (Fig. 4c, grey line). In the limit of *n* → ∞, Eq. 8 yields (Fig. 4c, blue line)

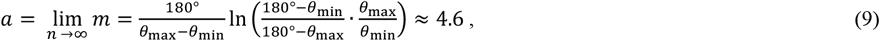

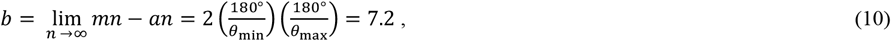

which is close to the parameters *a* = 5 and *b* = 8 in the AW law as given by Eq. 4 (Fig. 4c, black line), and within the range of parameter values inferred from biological data and the `LBIBCell` simulations (Fig. 2c). Aboav’s empirical values *a* = 5 and *b* = 8 are obtained with *θ*_min_ = 50.4° and *θ*_max_ = 160.7°, the lower bound of which is plausible only if trigons become distorted.

For small *n*, Eq. 8 predicts larger values for *m_n_* than what is observed. So, what happens for smaller *n*? Consider a regular triangle. To meet the angle constraint, it requires neighbouring polygons that each have 12 neighbours (Fig. 3a, purple). To maintain a lattice of equilateral polygons, this would require a tissue with a large variability in cell area (Eq. 3), and a strict pattern of large and small cells (Fig. 3a, purple). Such tissues are unlikely to emerge from biological processes. In the case of more physiological variabilities in apical cell areas, a triangular cell will border cells that have a smaller area than what is required for a regular dodecagon (*n*=12) according to the quadratic law (Eq. 3). As a consequence, such neighbouring dodecagonal cells would have a smaller side length than the triangular cell. To fit into a contiguous lattice, the polygons would then become stretched and their internal angles would no longer correspond to that of a regular polygon. Alternatively, the neighbours can assume a lower neighbour number than 12. Also in this case, the internal angles can no longer reflect that of a regular polygon as they would no longer add to 360°.

Accordingly, we extended the above formalism to allow for an irregular central polygon that deviates from the perfect internal angle *θ_n_* by an angle *θ*_i_ (Fig. 4d, inset; see Supplementary Information for details). In the limit of *n* → ∞, we recover the same value for the parameter *a* as for the regular polygon case (Eq. 9), but *b* now depends also on the maximal value of the deviatory angle *θ*_i_. We obtain an excellent fit to the AW law (*a*=5, *b*=8, Fig. 4d, black line) when we use *θ*_min_ = 360° − *θ_n_* − *θ*_i_ − *θ*_max_ and *θ*_max_ = 151.3°, and set the symmetric upper bounds of the angle mismatch to 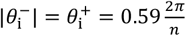, where 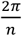 is the upper limit to maintain convexity (Fig. 4d, red line).

In fact, there is sufficient parametric freedom to reproduce all measured values of *a* and *b* (Fig. 2c) by adjusting the integration bounds for the angles, *θ*_min_ and *θ*_max_, and the upper bounds 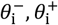 for the angle deviation *θ*_i_. Thus, the neighbour relationships in the *Drosophila* larval wing disc (*a*=4.76, *b*=8.34, grey) deviate slightly from the classical AW law (Eq. 4, Fig. 5a, black), but can be reproduced very well by the model (Fig. 5a, red line), and the predicted upper bound, 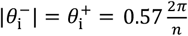, in the angle deviation matches that observed in the tissue very well (Fig. 5b). Importantly, even though we introduced the irregularity assumption to explain the neighbour relationships of cells with few neighbours, it applies to all polygon types. In agreement with this, the great majority of epithelial cells deviate from a regular shape as measured by their ellipticity, and the irregularity of the apical sides of epithelial cells does not dependent on the cells’ neighbour number (Fig. 5c).

**Figure 5:**
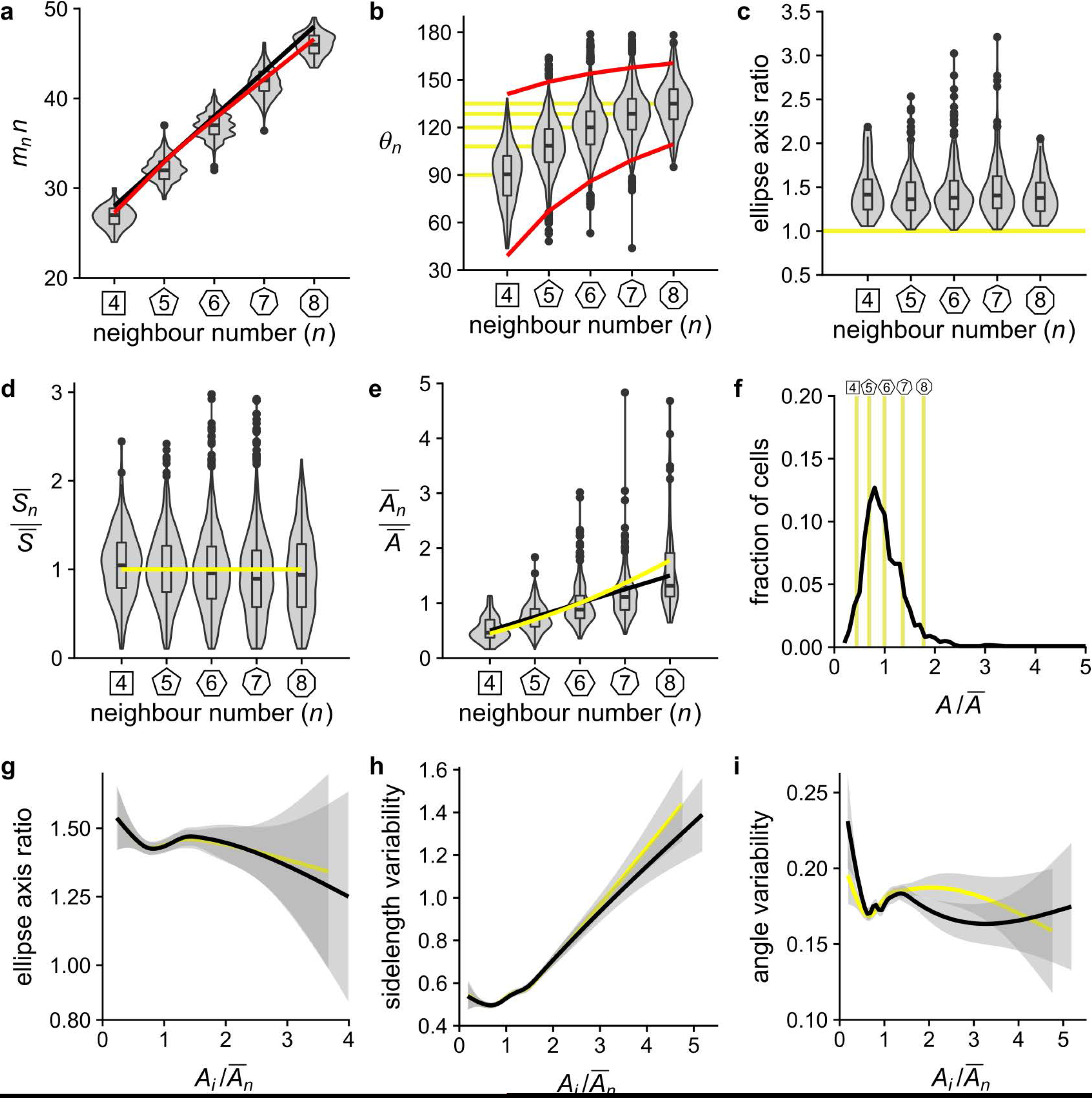
Individual apical areas in epithelia are irregular. **(a)** The product of *m_n_* · *n* versus neighbour number *n* for cells in the *Drosophila* wing disc follows the AW law on average (black line), and is approximated very well by the analytical approximation for irregular polygons (red line) when optimizing the angle bounds 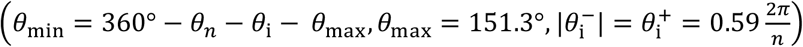 to match the AW law (*a*=4.76, *b*=8.34) that best matches the data (Fig. 2c). **(b)** Angle versus polygon type for cells in the *Drosophila* wing disc. The observed average angles are those of regular polygons (yellow lines). The red solid lines mark 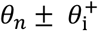, the predicted angle range of the irregular central polygon with 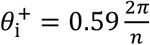. **(c)** Ellipticity of cells in the *Drosophila* wing disc as measured by the ratio of the axes of fitted ellipses. A ratio of 1 (yellow line) corresponds to regular polygons. **(d)** Normalised side length versus polygon type for cells in the *Drosophila* wing disc. The mean deviates from the ideal curve 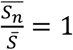 (yellow line). **(e)** The relative average cell area, 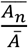, for cells in the *Drosophila* wing disc. The average is linearly related to the number of neighbours, *n*, in the *Drosophila* wing disc, and follows the linear Lewis’ law (Eq. 2, black line) rather than the quadratic law (Eq. 3, yellow line). **(f)** Normalised area distribution for cells in the *Drosophila* wing disc. The yellow vertical lines mark the areas for each polygon type required to form a regular polygon lattice (Eq. 3). **(g-i)** Ellipticity, as measured by the ratio of the axes of fitted ellipses, (f), variation of side length (g), and variation in the angles (h) of single cells in the *Drosophila* wing disc versus the fold-deviation of their apical area, *A_i_*, from that appropriate for their polygon type, 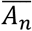, according to either the linear Lewis’ law (Eq. 2, black line) or the quadratic law (Eq. 3, yellow line). Variability in side length and angles was calculated for each cell as 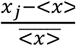, where *x_j_* refers to side length or angle, respectively, of each cell *j*, and < *x* > indicates the expected value for the particular polygon type of the cell.

Much as the internal angles, we expect the side lengths to deviate from that of a regular polygon. In a regular polygonal lattice, all side lengths should be equal (Eq. 3, Fig. 5d, yellow line), and the average normalised side lengths are indeed close to one (Fig. 5d). Single cells mostly deviate from these averages, much as only the average area per polygon type follows Lewis’ law (Eq. 2, Fig. 5e, black line). There are two sources for the observed irregularity. For one, the cellular processes result in a spatial distribution of apical areas in epithelia that are not consistent with the requirements of regular polygonal lattices (Fig. 3a). Secondly, apical areas in epithelia follow a continuous area distribution (Fig. 5f, black line) while the quadratic law (Eq. 3) specifies discrete optimal areas for each polygon type (Fig. 5f, yellow lines). With regard to the second point, we indeed observe that cell shape, side lengths, and angles deviate the least from that of a regular polygon when a cell’s neighbour number corresponds to the apical area according to the linear Lewis’ law (Eq. 2, Fig. 5g-i, black line) or the quadratic law (Eq. 3, Fig. 5g-i, yellow line).

## CONCLUSION

We conclude that the AW law, like Lewis’ law, emerges in epithelial tissues because cells minimise the overall lateral surface energy and thus the combined cell perimeter. By following the relationship between polygon area and polygon type as stipulated by Lewis’ law (Eq. 2) or the quadratic law (Eq. 3), the difference in side lengths is minimised between cells. By following the neighbour relationships that are described by the phenomenological AW law, the internal angles are closest to that of a regular polygon while adding to 360° at each vertex. Both the linear Lewis’ law and the AW law deviate from the curve predicted for a regular polygonal lattice. Regular polygonal lattices, however, require strict patterns of correctly sized cells. Given that cell growth and cell division processes result in a more variable spatial distribution of apical cell sizes, these cannot give rise to a regular polygonal lattice. The observed lattices therefore represent a trade-off to achieve the most regular cell shapes in a contiguous lattice with differently-sized apical areas.

## METHODS

### Image Processing

Cell boundaries and cell properties (cell area, polygon type, ellipse fit axes) were obtained in (Kokic et al., submitted). In addition, the side lengths and the internal angles of all cells were extracted using EpiTools [5] and the Fiji [17] plugin Tissue Analyzer [18]. To this end, the EpiTools source code was modified to also export the IDs of the *n_i_* neighbours of a cell to permit the calculation of *m_n_*. R scripts were used to extract and analyse data, calculate statistics, and plot the results.

### Set-Up of the LBIBCell simulations

The `LBIBCell` simulations were set up as described in detail before (Kokic et al, submitted) and is summarised in the Supplementary Material.

### Data and Software availability

All source codes are freely available at:

https://git.bsse.ethz.ch/iber/Publications/2019_vetter_kokic_aboav-weaire-law.

## Acknowledgements

We thank Davide Heller and Luis M. Escudero for providing access to their raw data. This work has been supported through an SNF Sinergia grant to DI, and grants BFU2012-34324 and BFU2015-66040 (MINECO, Spain) to FC.

## Author Contributions

DI, RV developed the theoretical framework, MK carried out all image processing, simulations and created all figures, HG, LH, BG contributed preliminary image processing, modelling and data analysis, AI, GV-F, FC contributed the data. DI, RV, FC, MK wrote the manuscript. All authors approved the final manuscript.

## Supplementary Material

### 1 Analytical approximation to Aboav—Weaire’s law

#### 1.1 Approximation for regular polygons

**Figure S1:**
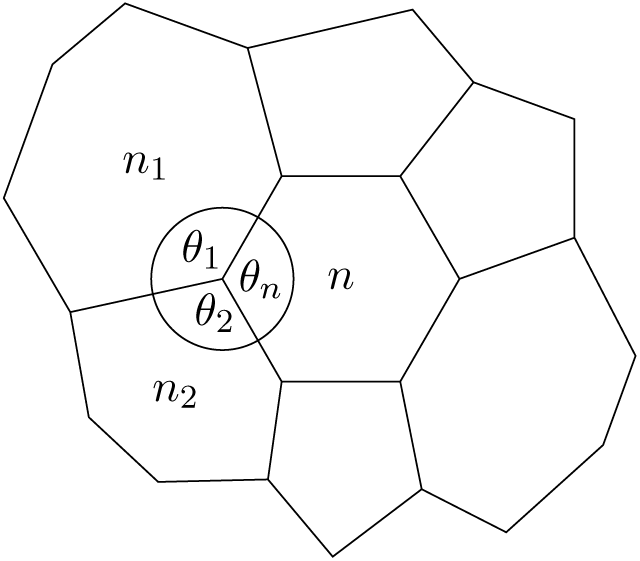
Illustration of a cell neighbourhood with regular central cell.

Consider the case of a regular polygon with *n* edges in the centre of a local epithelial cell neighbourhood. The interior angles of a regular polygon are given by

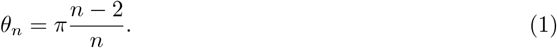

Inversely, the number of edges can be obtained from the interior angles as

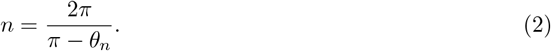

Assuming that all junctions are tri-cellular as illustrated in Fig. S1, and given *θ_n_*, the average angle left for the two neighbouring cells is

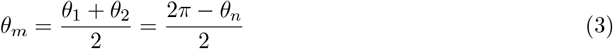

at each vertex. If *θ*_1_ = *θ*_2_ and also both neighbour cells are regular polygons, the mean neighbour number of all neighbours of a cell with *n* neighbours follows as

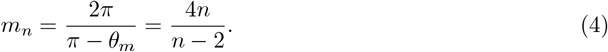

However, in most polygonal lattices, *θ*_1_ and *θ*_2_ differ. We are therefore now averaging over all allowed values of *θ*_1_:

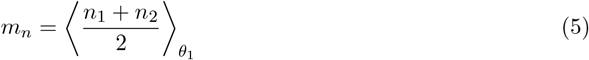

where

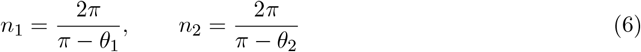

with

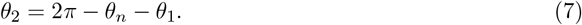

The average is taken over a physiological range [*θ*_min_, *θ*_max_] (to be specified later):

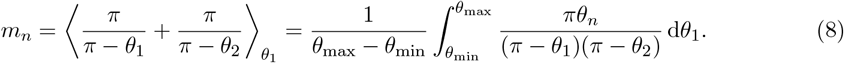

Notice that this definition is symmetric in *θ*_1_, *θ*_2_, i.e., one could also average over *θ*_2_ instead. The integral in Eq. 8 can be cast into the form

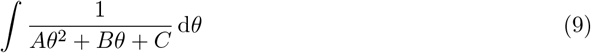

with coefficients

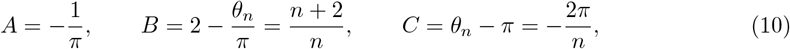

which evaluates to

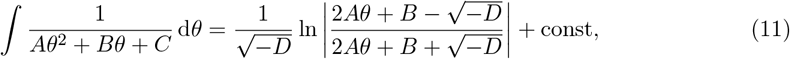

where

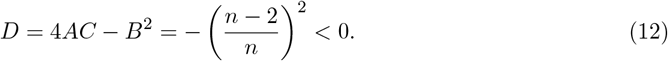

Therefore, after some basic algebra, Eq. 8 can be evaluated to

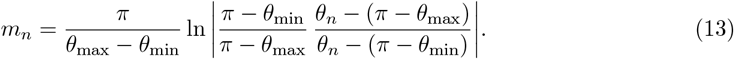

Eq. 13 requires careful selection of the integration bounds *θ*_min_, *θ*_max_, which can be functions of *n* in general. Trigons are the polygons with the lowest possible neighbour number, and assuming regular polygons, a consistent choice for the minimum allowable angle is

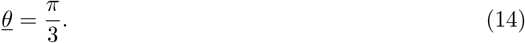

For the upper bound, a natural choice appears to be

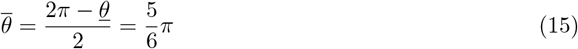

(corresponding to *n* = 12), as cells with more than 12 edges are hardly ever observable in epithelial tissue. Since this rationale applies to both angles *θ*_1_ and *θ*_2_ alike, and since they are coupled via Eq. 7, the bounds must be applied symmetrically, yielding

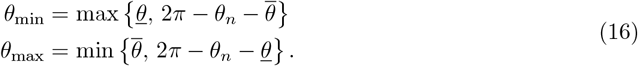

As long as *n* ≤ 12, this evaluates to *θ*_min_ = 7*π*/6 − *θ*_n_ and *θ*_max_ = 5*π*/6 (Fig. S2). With these integration bounds, Eq. 13 becomes

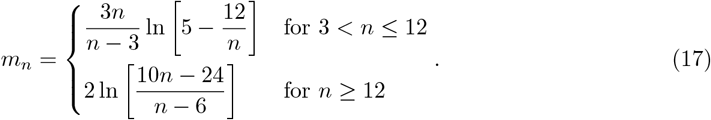

This relationship approximates Aboav–Weaire’s empirical law very well for *n* ≥ 6 (Fig. S3). Aboav–Weaire’s law can thus be explained for large *n* by the cell’s tendency toward a regular shape under the constraint that the three angles at cell junctions must sum up to 2*π*.

**Figure S2:**
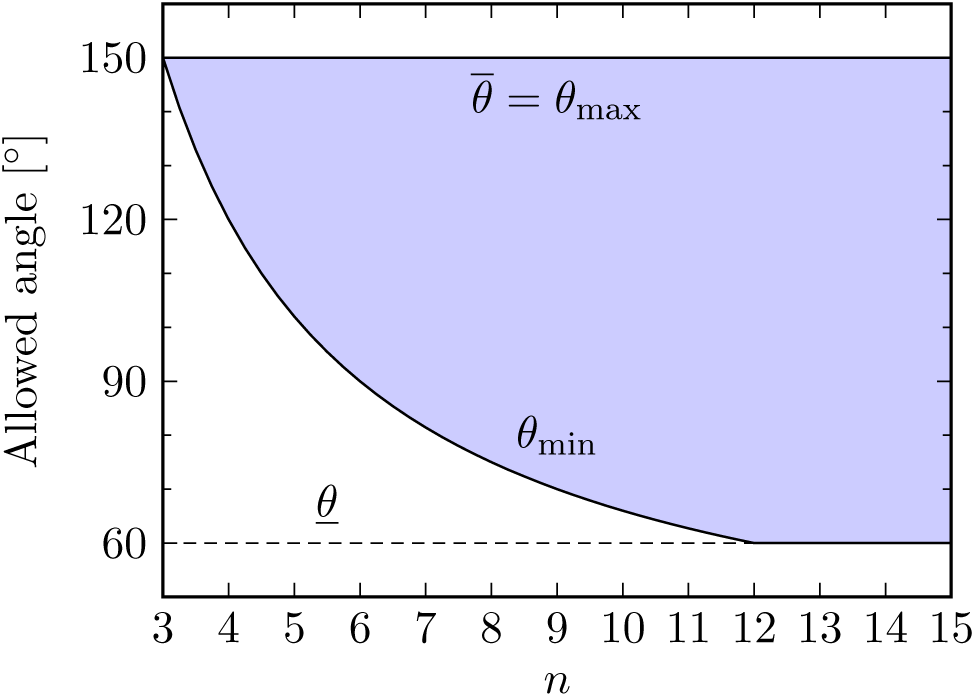
Integration bounds as defined in Eq. 16. The blue region indicates the integration domain.

**Figure S3:**
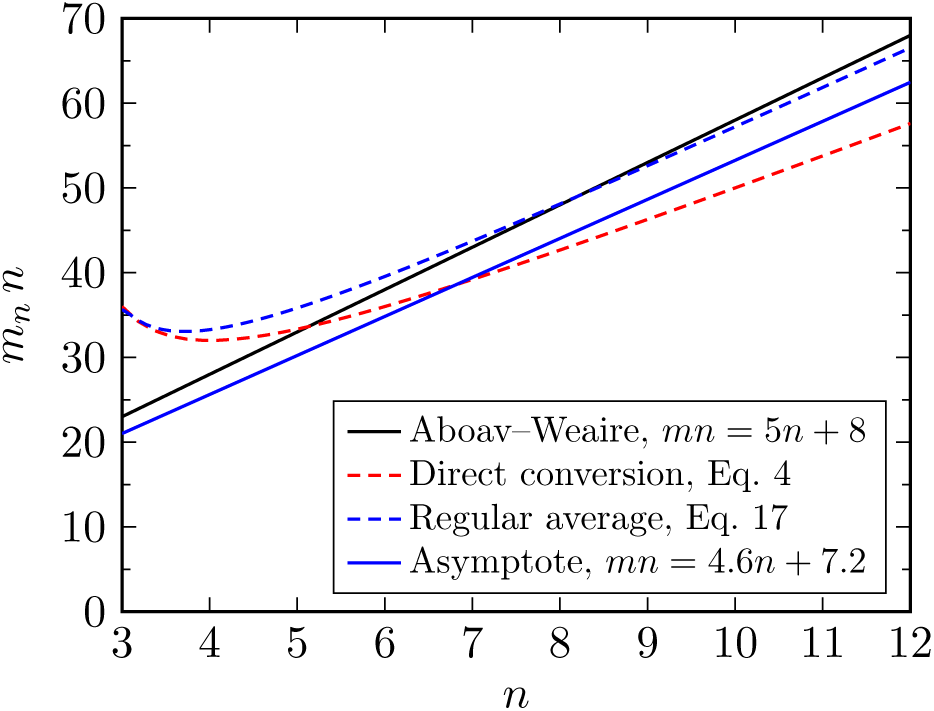
Comparison of analytical approximations to Aboav–Weaire’s law for regular polygons.

We now consider the asymptotic behavior in the limit of large *n*. Aboav’s empirical law reads

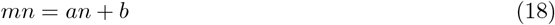

with coefficients *a* = 5, *b* = 8 originally observed in polycrystals [1]. Weaire derived the coefficients *a* = 5, *b* = 6 under a mean field approximation of polygonal regularity, which led him to suggest that *b* = 6 + Var(*n*) [17]. Taking the limit in Eq. 13, we find a linear asymptote with

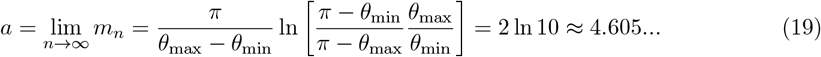

and

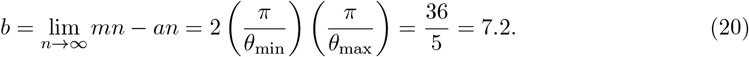

Both of these values lie remarkably well within the experimentally observed range of the coefficients for epithelial tissue (see Fig. 2c in the main article). Other values can be obtained if the physiological range over which the average is taken is chosen differently. The dependence of *a* and *b* on the integration bounds is shown in Fig. S4 along with the inverse relationship. As an example, Aboav’s empirical values *a* = 5, *b* = 8 are obtained if *θ*_min_ ≈ 50.4° and *θ*_max_ ≈ 160.7°, which are physiologically plausible values for epithelial tissue. Taking the mean epithelial coefficients *a* = 4.8, *b* = 8 (Fig. 2c) as a reference, we find the slightly tighter bounds *θ*_min_ ≈ 51.9° and *θ*_max_ ≈ 156.0°.

**Figure S4:**
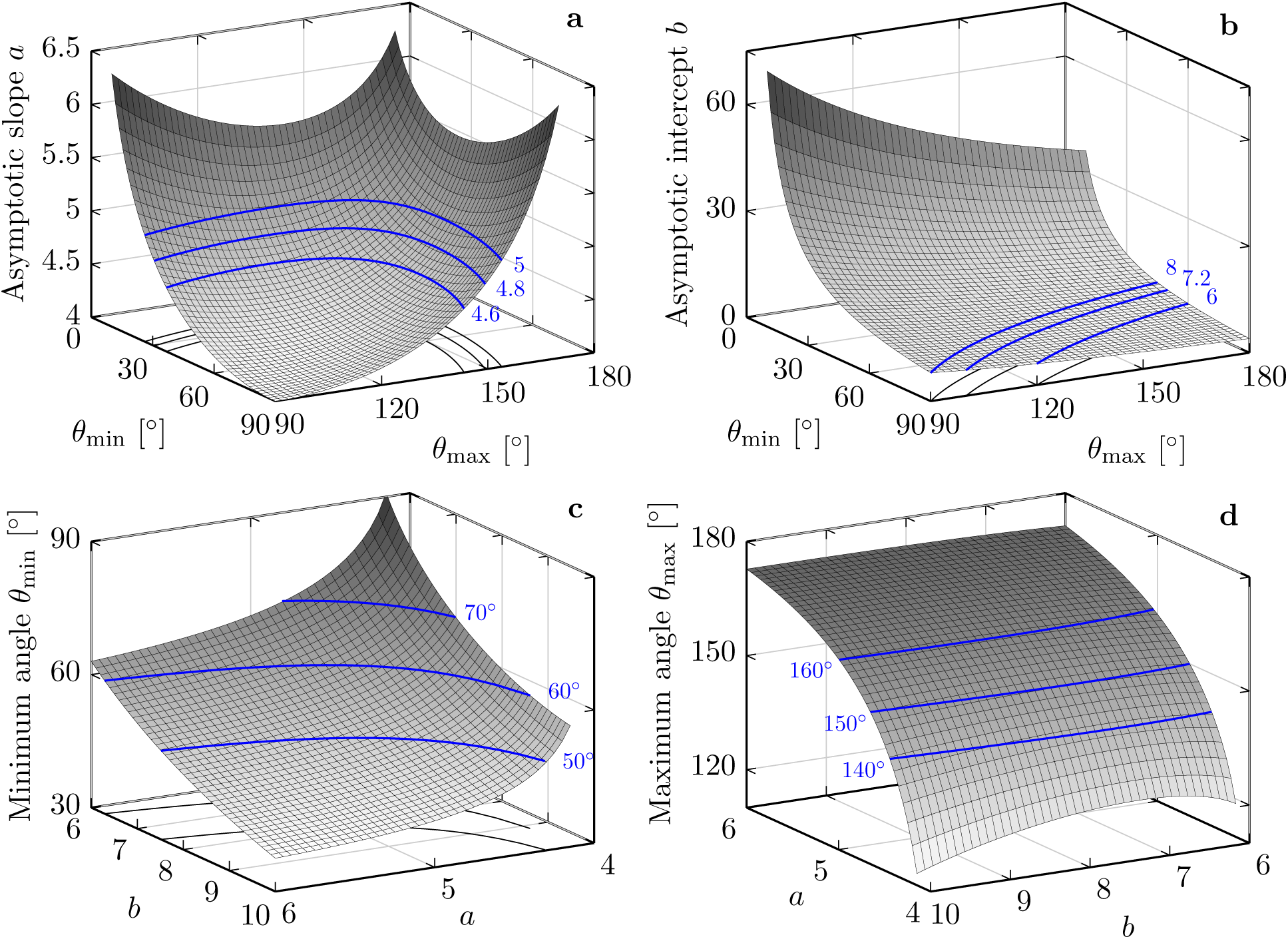
Asymptotic behavior of the analytical approximation in the limit *n* → ∞. (a-b) Line coefficients *a* and *b* as a function of the integration bounds *θ*_min_ and *θ*_max_. Typical values for *a* and *b* are highlighted as blue contour lines. Aboav–Weaire’s linear law can be obtained asymptotically by intersecting the contours for the desired values of a and b, yielding the physiological range for the angles. (c-d) Inverse relationship, obtained by numerical inversion of Eqs. 19 and 20.

#### 1.2 Generalisation to irregular polygons

**Figure S5:**
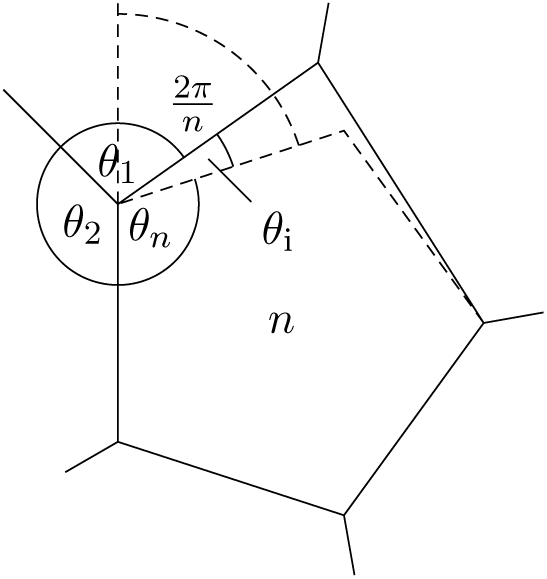
Illustration of the neighbourhood of an irregular polygon. The internal angle deviates from that of a regular polygon, *θ*_n_, by the angle *θ*_i_. The dashed lines mark the outline of a regular pentagon in the centre, as well as the maximal angle 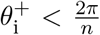 for *θ*_i_ to maintain convexity. The angles of the three polygons, *θ*_n_ + *θ*_i_, *θ*_1_, *θ*_2_, must add to 2*π*.

Evidently, the regularity assumption on the polygonal cell shape breaks down for small *n*. Eq. 17 predicts higher polygon types around triangles, quadrilaterals and pentagons than can be observed in many cellular arrangements (Fig. S3). In real epithelial tissue, cells with large area are not available discretionarily, as required for the regularity assumption to hold for small *n*. Additionally, side lengths between neighbouring cells must match at shared edges, posing further limitations on the attainable cell regularity in a contiguous lattice. Therefore, a generalisation of the above formalism is proposed here for irregular central polygons.

We now consider a deviation *θ*_i_ from the internal angle of a regular polygon (Eq. 1) (Fig. S5), and calculate the expected neighbour number of direct neighbours by averaging over a physiological range 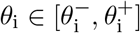, yielding

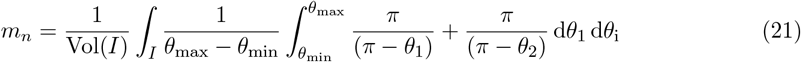

subject to the modified angle sum constraint

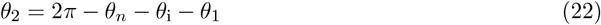

and modified integration bounds

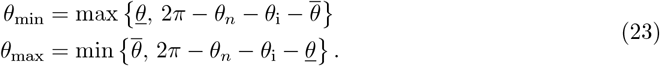

Here, *I* denotes the set of possible irregularity angles

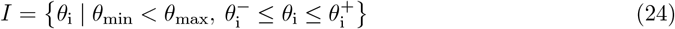

which depends on *n* through Eqs. 23 and 1, and

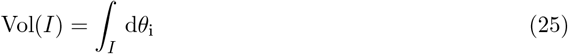

is its volume (in the present one-dimensional case, its length). A natural choice for the permitted degree of irregularity appears to be to set the bounds 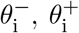 to a certain fraction of the turning angle of regular polygons, *π* − *θ_n_* = 2*π*/*n*, which represents the maximum excess angle that will maintain convexity of the polygon (Fig. S5). It is therefore reasonable to set

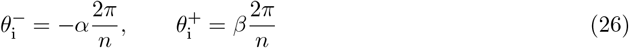

with free (but possibly interrelated) coefficients *α, β* ∈ [0, 1).

**Figure S6:**
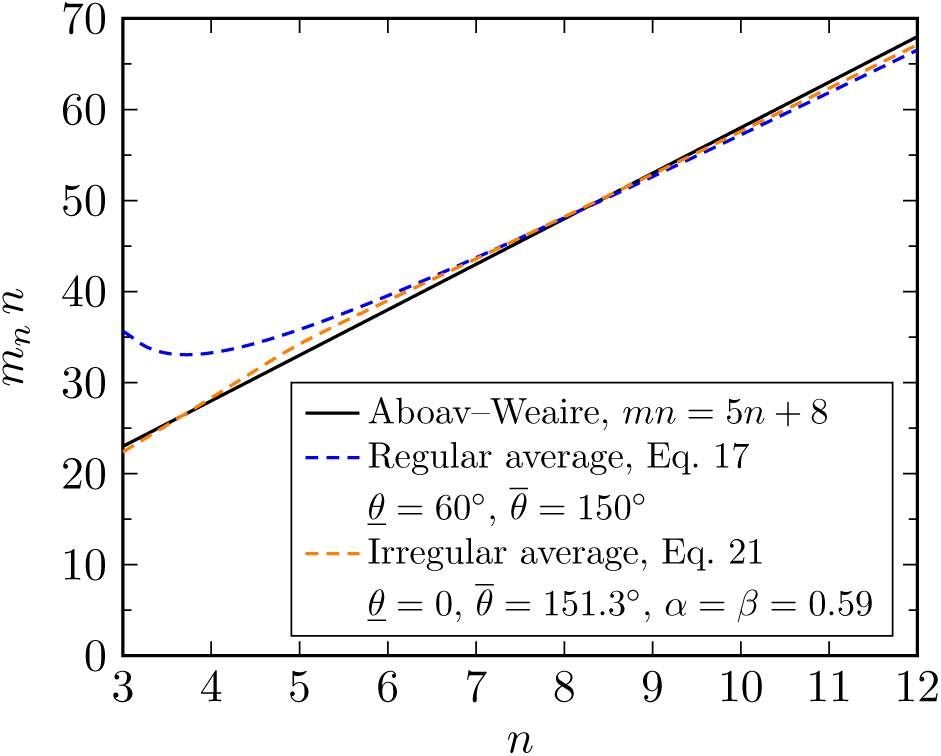
Comparison of analytical approximations to Aboav–Weaire’s law for irregular polygons.

The inner integral in Eq. 21 can be evaluated using Eq. 13, yielding

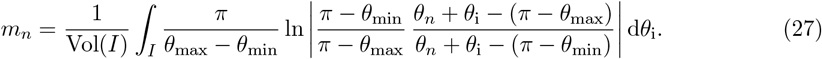

However, Eq. 27 is a non-elementary integral as it can be expressed in terms of dilogarithm functions [7]. Taking also the complicated integration domain *I* into account, a closed-form solution appears to be inaccessible. Nevertheless, with the assumptions used for the irregularity bounds in Eq. 26, it is easy to recognize that the same asymptotic slope *a* as for the regular case is recovered, since Eq. 27 converges to Eq. 13 as 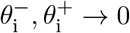, which is the case for *n* → ∞. The asymptotic intercept *b*, on the other hand, depends on the granted range of irregularity.

In order to calculate *m_n_* for given *n*, we evaluate the double integral numerically using the acceptance-rejection method with uniform sampling in the domain 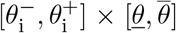 in MATLAB 2018a. The integral offers sufficient parametric freedom to get very close to the linear Aboav–Weaire relationship over the entire range of polygon types (Fig. S6), and it still does if 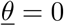 is fixed, such that the lower integration bound *θ*_min_ is essentially governed by the symmetry requirement of *θ*_1_ and *θ*_2_. Furthermore, restricting the irregularity angles to symmetric domains by setting *α* = *β* leaves us with two free parameters, 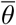 and *β*. A curve fit of our model to the empirical line *mn* = *an+b* with *a* = 5 and *b* = 8 yields least squared errors for 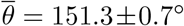 and *β* = 0.59±0.042 with an adjusted coefficient of determination of 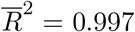 (Fig. S6). Taking the mean epithelial coefficients *a* = 4.8, *b* = 8 (see Fig. 2c in the main article) as a reference, we find 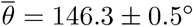 and *β* = 0.52 ± 0.03 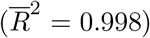, both of which are physiologically plausible values for epithelial tissue.

In summary, the linear Aboav–Weaire law emerges naturally for geometric reasons in contiguous, two-dimensional cellular lattices when a limitation in locally available cell areas and matching side lengths enforce polygonal irregularity, i.e., interior angles deviating from those of regular polygons. For high-*n* polygons (*n* > 6), it can even be explained by the cells’ tendency toward regular polygonal shapes alone. We expect this explanation of Aboav–Weaire’s law to be applicable also to other systems partially driven by surface minimisation and area variability, such as soap bubble froths or polycrystals. Our theoretical results might also be appreciable in terms of contact graph theory, of which the present polygonal space-filling packing represents a special case.

### 2 LBIBCell Simulations

#### Simulation framework

We used the software `LBIBCell` [15] to simulate the detailed epithelial tissue dynamics. `LBIBCell` simulates a 2D plane of the epithelial tissue, and thus allows us to simulate the apical cell dynamics, or more precisely a plane where cells adhere laterally (Fig. 2a, main manuscript). `LBIBCell` represents the extracellular and cytoplasmic spaces of the tissue as Newtonian fluids and incorporates the fluid-structure interaction with the elastic cell boundaries via an immersed boundary method [10]. The Navier-Stokes equation for the fluid dynamics is solved using the Lattice Boltzmann method, i.e., the fluid and its motion are realised by particle ensembles which stream and collide on a 2D regular lattice [2]. Cell boundaries are geometrically highly resolved and cell boundaries and cell-cell junctions are modelled with elastic springs.

#### LBIBCell Simulations for Figure 2

The proliferating *Drosophila* larval wing disc tissue was simulated as described in (Kokic et al, submitted); the parameter values are summarised in Table S1. Simulations with two different cell division rules were carried out 10 times each. According to the first cell division rule, Hertwig’s rule [5,3,8], cells divide perpendicular to their longest axis. In the second set of simulations, the orientation of the cell division axis was randomised. Details on the implementation of the cell division rules can be found in [15].

#### LBIBCell Simulations for Figure 3

To generate polygonal lattices of regular polygons, centres of circular cells were placed at sites corresponding to the centres of regular polygons in different Euclidean tilings by convex regular polygons (i.e., {6, 6, 6}, {4, 8, 8}, {4, 6,12}, {3,12,12}). The relative sizes of the cells followed the quadratic law (Eq. 3 in the main manuscript). In a second step, the circular cells were enlarged uniformly until they collided at single points with their neighbours. These start structures (`LBIBCell` input) were created using Inkscape [6], Fiji [13,14] and R [16]. A detailed description of the pipeline and scripts are available on https://git.bsse.ethz.ch/iber/Publications/2019_vetter_kokic_aboav-weaire-law. In a last step, cells were grown uniformly until the entire lattice was filled with cells and cells thus formed a contiguous lattice. In order to prevent gaps, the cell junction search radius was increased to its maximum value (cf. Figure S3 in Kokic et al., submitted; Table 1), and the amount of mass that was added intracellularly was subtracted from the extracellular fluid (cf. `Cell Junction` and `Membrane Channels` in Table S1). All other `LBIBCell` parameter values were set to the same values as for the *Drosophila* larval wing disc (Table S1), and all subsequent quantifications were carried out as described in (Kokic et al., submitted).

### 3 Supplementary Tables

**Table S1:** Parameter values for the simulation of *Drosophila* larval wing disc epithelia. Parameters in grey-highlighted rows only affect the numerical accuracy of the simulations. The other parameter values are specific to the biological problem of interest, and were determined based on published quantitative data for the *Drosophila* larval wing disc pouch (Kokic et al, submitted). This table is reproduced with modifications from (Kokic et al, submitted).

|  |  |  |  |  |
| --- | --- | --- | --- | --- |
| **LBSolver** | Parameter Description | Value [LB units] | Value [SI units] | Determination / Literature |
| | $\tau_{\text{fluid}}$ - relaxation time | 2 | N/A | [11] |
| **BioSolver** | Parameter Description | Value [LB units] | Value [SI units] | Determination / Literature |
| **Cell Growth** | frequency of applying BioSolver | 1:1 | N/A | (Kokic et al, submitted) |
| | $I$ - total number of iterations | 64300 / 150000 | 24-96h AEL / 0-168h AEL | (Kokic et al, submitted) |
| | $b$ - Gompertz constant | 11.331 | 11.331 | [9] |
| | $\tau$ - Gompertz constant | 0.8555 | 143.7773h/168h | [9] |
| **Membrane Tension** | frequency of applying BioSolver | 1:10 | N/A | (Kokic et al, submitted) |
|  | Hookean spring constant | 0.01 | N/A | [4] |
| **Cell Junction** | frequency of applying BioSolver | 1:10 | N/A | (Kokic et al, submitted) |
|  | search radius for junction building | 1.0 | N/A |  |
|  | length of intercellular cell junction | 0.3 | 27nm | 16-39nm [12] |
|  | Hookean spring constant | 0.1 | N/A | [4] |
| **Cell Division** | frequency of applying BioSolver | 1:1000 | N/A | (Kokic et al, submitted) |
| | maximal cell area | 1771 | $14.1\mu\text{m}^2$ | [4] |
| | mean cell division threshold | 837 | $7\mu\text{m}^2$ | [4] |
| | std cell division threshold | 256 | $2\mu\text{m}^2$ | [4] |
| **Cell Extrusion** | frequency of applying BioSolver | 1:100 | N/A | (Kokic et al, submitted) |
| | minimal cell area | 100 | $0.8\mu\text{m}^2$ | [15, 4] |
| **Membrane Channels** | frequency of applying BioSolver | 1:1 | N/A | [15] |
|  | growth source | 0.01 | N/A | [15] |

